# Comparative Analysis of the Effects of PSPH and PHGDH Inhibitors on Tumor Cell Proliferation

**DOI:** 10.1101/2025.04.28.650903

**Authors:** Yanbing Wang, Longze Sha

**Affiliations:** State Key Laboratory of Common Mechanism Research for Major Diseases, Department of Biochemistry and Molecular Biology, Institute of Basic Medical Sciences, Chinese Academy of Medical Sciences and Peking Union Medical College, Beijing 100005, China; State Key Laboratory of Complex, Severe and Rare Diseases, Peking Union Medical College Hospital, Peking Union Medical College and Chinese Academy of Medical Sciences, Beijing, 100730, China; Neuroscience Center, Chinese Academy of Medical Sciences, Beijing, 100005, China

**Keywords:** Tumor metabolism, Serine biosynthesis, PHGDH, PSPH, α-Ketoglutarate, Cell proliferation

## Abstract

Serine metabolism plays a critical role in supporting the rapid proliferation of tumor cells, with PHGDH recognized as a key rate-limiting enzyme and therapeutic target. However, whether its antitumor effects are solely dependent on serine metabolism remains under debate. In this study, we systematically compared the effects of PHGDH versus PSPH inhibitors on intracellular serine levels and tumor cell proliferation. Although multiple PSPH inhibitors significantly reduced intracellular serine concentrations, they failed to effectively inhibit cell proliferation. In contrast, PHGDH inhibitors exhibited robust antiproliferative activity under both serine-deprived and serine-replenished conditions. Further supplementation with α-ketoglutarate, a downstream metabolite of PHGDH, partially reversed this inhibitory effect. These findings suggest that the antitumor activity of PHGDH inhibitors is not solely attributable to serine metabolism inhibition, but rather results from the coordinated disruption of multiple metabolic pathways, offering a novel perspective for metabolism-targeted cancer therapy.

## 1. Introduction

Cancer remains the second leading cause of death worldwide, with an estimated 20 million new cases diagnosed globally in 2024 and approximately 9.7 million cancer-related deaths[1]. During malignant progression, cancer cells acquire a series of adaptive traits collectively termed the “hallmarks of cancer,” including uncontrolled proliferation, enhanced metastatic capacity, and evasion of cell death[2]. These traits are sustained by metabolic reprogramming, which supports the high bioenergetic and biosynthetic demands associated with rapid growth[3]. Among these reprogrammed pathways, serine metabolism plays a critical anabolic role in processes such as protein synthesis, nucleotide production, and antioxidant defense, and is broadly recognized as essential for highly proliferative cells[4, 5].

Intracellular serine is derived through two main routes: uptake from the extracellular environment via specific transporters, and de novo biosynthesis[6]. De novo serine biosynthesis is a tightly regulated, three-step enzymatic process. First, 3-phosphoglycerate (3PG), a glycolytic intermediate, is converted to 3-phosphohydroxypyruvate (3PHP) by phosphoglycerate dehydrogenase (PHGDH). Second, 3PHP undergoes a transamination reaction catalyzed by phosphoserine aminotransferase 1 (PSAT1), forming phosphoserine (p-Ser) by utilizing glutamate as a nitrogen donor and generating α-ketoglutarate (α-KG) as a by-product. Finally, phosphoserine phosphatase (PSPH) catalyzes the dephosphorylation of p-Ser to produce serine[7]. Recent studies have identified PHGDH as a key rate-limiting enzyme in this pathway, with its expression markedly upregulated in various cancers, including approximately 40% of breast cancers and 50–80% of lung cancers[8-10], thus positioning PHGDH as a promising therapeutic target.While PHGDH inhibition has been shown to markedly reduce tumor cell growth in some contexts, whether this effect is primarily mediated through suppression of serine synthesis remains contentious[11]. PHGDH is also involved in several other critical metabolic processes, including one-carbon metabolism, redox homeostasis via NAD^+^/NADH balance, α-KG production, and nucleotide biosynthesis[12]. Notably, studies have demonstrated that even in the presence of sufficient extracellular serine, inhibition of PHGDH can still suppress cell proliferation, suggesting that the role of this pathway extends beyond simply providing serine[8, 13].

In this study, we aimed to determine whether the tumor-suppressive effects of PHGDH inhibition are entirely dependent on serine metabolism. By developing new PSPH inhibitors, we found that its inhibition alone did not significantly suppress tumor cell proliferation,though the levels of serine is decreased. Moreover, supplementation with exogenous serine failed to fully reverse the antiproliferative effects of PHGDH inhibitors, indicating that PHGDH inhibition exerts broader metabolic consequences beyond serine depletion. We found that supplementation with α-ketoglutarate partially reversed the antiproliferative effect of PHGDH inhibitors, further supporting the notion that PHGDH suppresses tumor growth through multiple metabolic pathways beyond serine biosynthesis.These findings offer a revised understanding of PHGDH’s role in tumor metabolism and highlight the multifaceted consequences of its inhibition. Our results not only deepen the mechanistic insight into metabolic vulnerabilities in cancer but also inform future directions for drug development and target selection in metabolic-based cancer therapies.

## 2. Materials and methods

### 2.1 Surface Plasmon Resonance

Surface plasmon resonance (SPR) measurements were conducted using a Biacore system. The sensor chip (CM5, Cytiva) was activated using 1-ethyl-3-(3-dimethylaminopropyl) carbodiimide (EDC, Cytiva) and N-hydroxysuccinimide (NHS, Cytiva) at a flow rate of 10 μL/min. Target proteins were immobilized onto the chip surface via amine coupling chemistry, followed by blocking of unreacted sites with ethanolamine at the same flow rate.Test compounds were prepared in a 96-well plate and serially diluted to various concentrations (0.3125–10 μM). The diluted compounds were injected in ascending concentration order at a flow rate of 30 μL/min for 150 seconds. After each injection, the chip surface was regenerated with 10 mM glycine-HCl (pH 2.0) for 5 minutes. This cycle was repeated until all concentrations of each compound were tested.Sensorgrams were globally fitted to a 1:1 Langmuir binding model using Biacore Insight Evaluation Software (Cytiva, Marlborough, MA, USA) to determine the association (ka) and dissociation (kd) rate constants, as well as the equilibrium dissociation constant (KD).

### 2.2 Targeted metabolomics

We performed 600MRM analysis (Biotree) with LC–tandem MS (LC-MS/MS). Primary astrocytes were harvested by adding 1500 μL of acetonitrile-methanol-H2O (2:2:1, containing isotope internal standards) into an Eppendorf tube. The samples were then frozen in liquid nitrogen and thawed in a 37 °C water bath. The freeze-thaw cycle was repeated 3 times, after which the samples were vortexed for 30 s. After 15 min of sonication in an ice-water bath, the samples were incubated at -40 °C for 2 h. Then, the samples were centrifuged at 1000 g and 4 °C for 15 min. A total of 1200 μL of the supernatant from each sample was transferred to a new tube and dried with a centrifugal concentrator. Next, 120 μL of 60% acetonitrile was added to the Eppendorf tube to reconstitute the dried samples. The Eppendorf tube was vortexed until the powder was dissolved, followed by centrifugation at 1000 g and 4 °C for 15 min. Finally, 60–70 μL supernatant of each sample was transferred to a glass vial for LC-MS/MS analysis. A mixture of standard metabolites was prepared as the QC sample. LC separation was carried out with a UPLC System (1290, Agilent) equipped with a Waters Atlantis Premier BEH Z-HILIC column (1.7 μm, 2.1 mm × 150 mm). The mobile phase A was mixed H2O and acetonitrile(9:1), containing 10 mmol/L ammonium formate, and the mobile phase B was mixed H2O and acetonitrile (1:9) containing 10 mmol/L ammonium formate. The autosampler temperature was set at 4 °C and the injection volume was 1 μL. AB Sciex QTrap 6500 plus mass spectrometer was applied for assay development. Typical ion source parameters were as follows: IonSpray Voltage: +5500V/-4500V, Curtain Gas: 35 psi, Temperature: 400 °C, Ion Source Gas 1: 50 psi, Ion Source Gas 2: 50 psi. Raw data files generated by LC-MS/MS were processed with SCIEX Analyst Work Station Software (1.7.3), metabolites quantification was analyzed with BIOTREE BioBud(2.0.3).

### 2.3 Cell lines and primary astrocyte cultures

293T cells (for rPSPH production) and HCC70,BT-20 cells (for CCK8 assays) were grown in DMEM (CM15019, Macgene). All media were supplemented with 10% fetal bovine serum (FBS, 10099141, Thermo Fisher), 100 U/mL penicillin, and 100 mg/mL streptomycin (CC004, Macgene). The cerebral cortices of unsexed P0-P2 newborn KM pups were used for primary astrocyte culture. Specifically, the cerebral cortices were dissected and mechanically dissociated by repeated pipetting with a 1 mL plastic pipette and a 25-gauge needle. The cells were then resuspended in DMEM (CM15019, Macgene) supplemented with 20% FBS (10099141, Gibco), 100 U/mL penicillin and 100 mg/mL streptomycin (CC004, Macgene) and plated onto poly-d-lysine-coated (A3890401, Thermo Fisher) T75 cm2 flasks. After 7 days of growth, microglia and oligodendrocytes were removed following shaking overnight at 37 °C. Astrocytes were replated onto dishes coated with poly-D-lysine for subsequent experiments. Customized serine/glycine-free DMEM medium was ordered from Macgene.

### 2.4 Plasmids and purification of recombinant PSPH (rPSPH)

The human PSPH (P78330) DNA sequence was optimized, synthesized, and subcloned into the pcDNA3.1 vector to create the pcDNA3.1-PSPH-FLAG construct (Tsingke Biotech). After 48 hours, cells were lysed on ice using Cellytic lysis buffer (C2978, Sigma-Aldrich) containing a protease inhibitor cocktail (B14002, Bimake). Recombinant PSPH was then isolated through immunoprecipitation with an anti-FLAG M2 affinity gel (A2220, Sigma-Aldrich) and subsequently eluted from the beads using 3×FLAG peptide (F4799, Sigma). The purity and structural integrity of the purified proteins were evaluated using Coomassie blue staining.

### 2.5 CCK-8 assay

Cell viability was assessed using the CCK-8 assay (HY-K0301, MCE). HCC-70 and BT-20 cells were seeded into 96-well plates at a density of 5,000 cells per well. After cell attachment, the culture medium was replaced with fresh medium containing either PSPH or PHGDH inhibitors at final concentrations of 40 or 20, 10, 5, 2.5, 1.25, 0.625, 0.3125, 0.15625 or 0.07812 μM. After 4 days of treatment, 10 μL of CCK-8 solution was added to each well and incubated at 37 °C for 2 hours. Absorbance was then measured at 452 nm. All experiments were performed in triplicate.

### 2.6 Statistical analysis

Statistical analysis was performed with Prism 8.0 software (GraphPad). To assess differences between two experimental groups, unpaired two-tailed Student’s t tests were used for data that were normally distributed according to the Kolmogorov-Smirnov test. Dose–response curves were generated using nonlinear four-parameter logistic regression to calculate IC_50_ values. The data are presented as the means ± SDs or individual points. A p-value less than 0.05 was considered to indicate statistical significance.

## 3. Result

### 3.1 Identification and characterization of potent small-molecule inhibitors targeting PSPH

To elucidate the critical role of serine metabolism in tumor metabolic regulation, we focused on targeting phosphoserine phosphatase (PSPH), the terminal rate-limiting enzyme in the de novo serine biosynthesis pathway. According to our recently accepted work (Sha et al., accepted by Nature Chemical Biology), PSPH inhibition markedly blocks the dephosphorylation of O-phospho-L-serine to L-serine, without affecting upstream metabolic pathways, thereby offering high specificity. This makes PSPH an ideal target for directly assessing the contribution of serine metabolism to tumor growth and progression.We identified four small-molecule inhibitors, Z218484536 (in press), Z1444603284 (Fig. 1a), Z997780042 (Fig. 1d), and Z1444669980 (Fig. 1g), that effectively suppressed PSPH-mediated L-serine production in a dose-dependent manner. In our previous work, we identified the binding characteristics between PSPH and Z218484536. In this study, we further investigated the binding features of three additional PSPH inhibitors using molecular docking and surface plasmon resonance (SPR) analysis. Molecular docking revealed that Z1444603284 forms stable hydrogen bonds and electrostatic interactions with several key amino acid residues in the active site of PSPH, including Asp22, Lys158, Thr182, and Gly110. Additionally, its aromatic groups interact with hydrophobic residues such as Phe58 and Ala51, enhancing the stability and affinity of the complex (Fig. 1b). Similarly, Z997780042 establishes multi-point hydrogen bonds and electrostatic interactions with PSPH, targeting critical residues such as Lys158, Asp179, Asp22, and Arg65. Hydrophobic residues like Phe57, Val56, and Ala55 create a hydrophobic environment that stabilizes binding (Fig. 1e). Z14444669980 exhibits excellent compatibility with the PSPH active pocket, forming multiple hydrogen bonds with residues Asp22, Lys158, Ala51, Ser109, and Gly110, as well as hydrophobic and π-π interactions with Gly157 and Phe58, demonstrating strong binding affinity (Fig. 1d).

**Figure 1.**
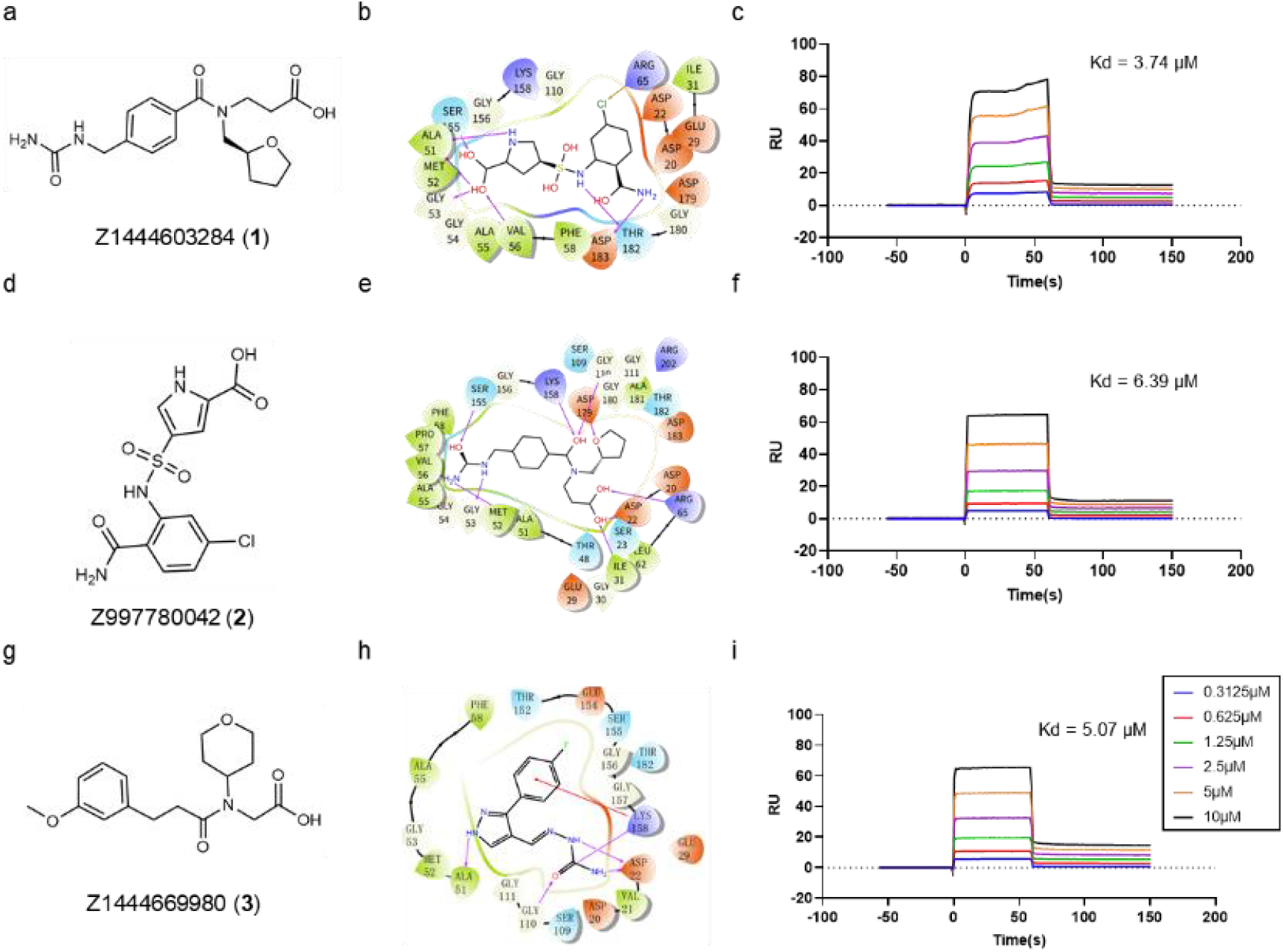
Identification and characterization of potent small-molecule inhibitors targeting PSPH. (a, d, g) Chemical structure of Z1444603284, Z997780042 and Z14444669980. (b, e, h) Molecular docking diagrams of three candidate compounds with PSPH active pockets, showing hydrogen bonding, electrostatic and hydrophobic interactions with multiple key amino acid residues. (c, f, i) Surface plasmon resonance (SPR) binding curves of three compounds with PSPH, The dissociation constants (Kd) of Z1444603284 (c), Z14444669980 (i) and Z997780042 (f) are 3.74 μM, 5.07 μM and 6.39 μM, respectively.

To confirm the binding affinity of these novel PSPH inhibitors, surface plasmon resonance (SPR) technology was employed to measure the three compounds to PSPH. The fitted sensorgrams indicated that all three compounds conform to the 1:1 binding model, yielding corresponding dissociation constants (Kd). Z1444603284 exhibited the highest affinity, with a Kd of 3.74 μM (Fig. 1c), followed by Z14444669980 with a Kd of 5.07 μM (Fig. 1f). Z997780042 showed relatively lower affinity, with a Kd of 6.39 μM (Fig. 1i). In summary, Z1444603284, Z997780042, and Z14444669980 specifically target PSPH and exhibit moderate binding affinity.

### 3.2 Cell-Type Specific Effects of PSPH and PHGDH Inhibitors on Serine Levels

To further clarify the efficiency of PSPH and PHGDH inhibitors in regulating serine metabolism, we employed four PSPH inhibitors, as well as two classical PHGDH inhibitors, Compound 18[14] and NCT-503[15]. These compounds were applied to primary cultured astrocytes and two tumor cell lines, HCC-70 and BT-20. Cells were cultured in DMEM medium lacking serine and glycine for 48 hours, after which they were harvested for targeted metabolomics analysis to determine intracellular L-serine levels. The results revealed significant differences in serine inhibition efficiency among different inhibitors across various cell types.

Z218484536 markedly reduced serine levels in primary astrocytes but showed no significant effect in HCC-70 and BT-20 tumor cells (Fig. 2a).These cell-type-specific differences suggest that PSPH may be regulated by post-translational modifications (PTMs). Alterations in PTMs, which are common in cancer cells, could affect the conformation, stability, or binding affinity of PSPH to small-molecule inhibitors[16]. In contrast, Z1444603284 and Z997780042 effectively reduced serine levels in both tumor cell lines but had minimal effects on astrocytes (Fig. 2b-c). These differences may be attributed to the distinct structural features of the compounds and their specific binding affinity to the PSPH active site. Additionally, Z1444669980 (Fig. 2d), NCT-503 (Fig. 2e), and Compound 18 (Fig. 2f) demonstrated differential degrees of serine inhibition efficacy across all three cell types.

**Figure 2.**
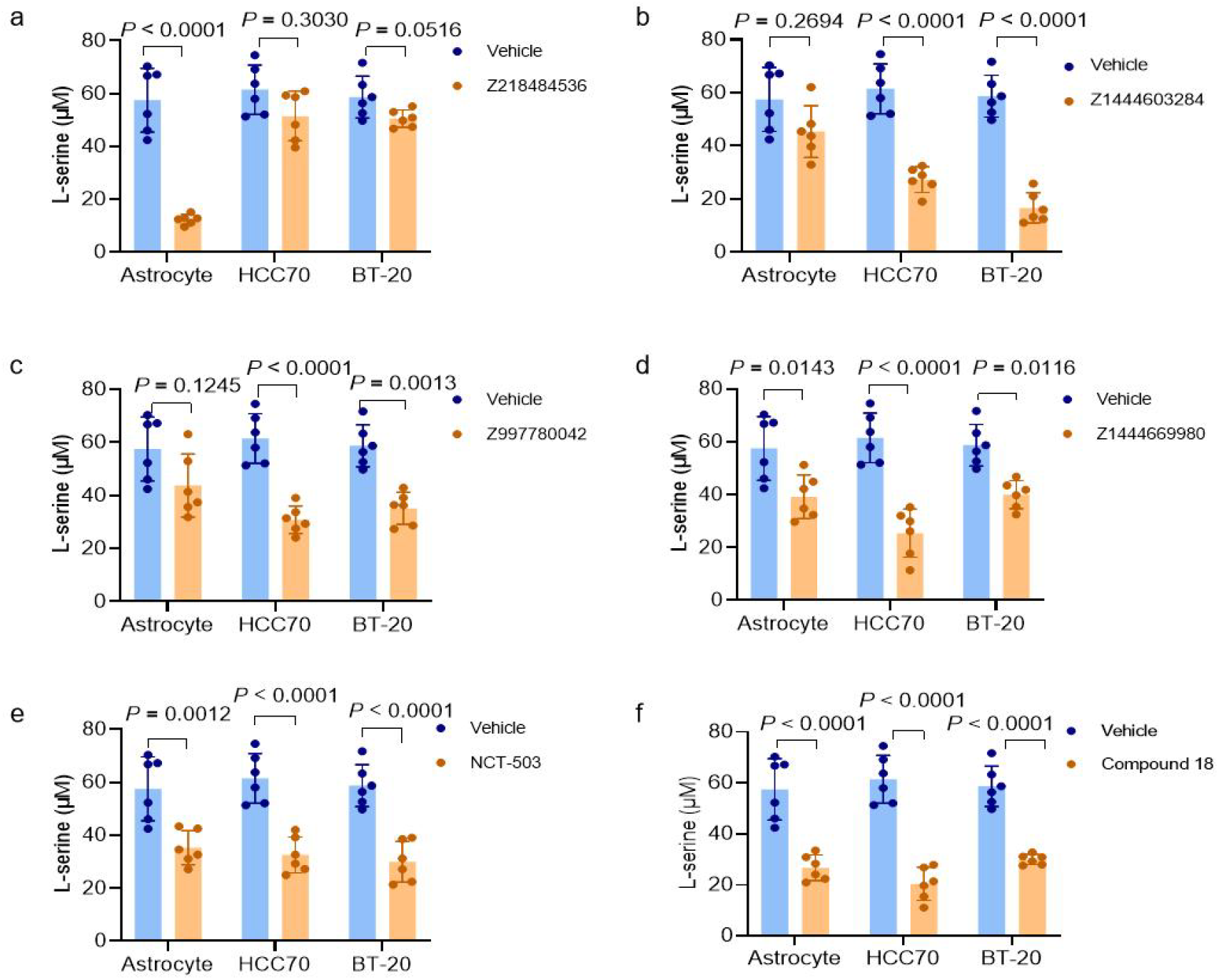
Cell-Type Specific Effects of PSPH and PHGDH Inhibitors on Serine Levels. Primary astrocytes and tumor cell lines HCC70 and BT20 were cultured in DMEM medium lacking serine and glycine and treated with four PSPH inhibitors Z218484536 (a), Z1444603284 (b), Z997780042 (c), and Z1444669980 (d) at a final concentration of 4 μM, as well as two PHGDH inhibitors NCT503 (e) and Compound 18 (f) at a final concentration of 2 μM for 48 hours, followed by targeted metabolomics analysis to measure intracellular L-serine levels. Data are presented as mean ± standard deviation (SD), with each dot representing an individual biological replicate (n = 6). Statistical analysis was performed with two-tailed Student’s t test (a-f).

In summary, we identified and validated several inhibitors of serine metabolism enzymes. Their distinct inhibitory profiles across cell types underscore the complexity of serine metabolic regulation. However, whether suppression of serine biosynthesis alone is sufficient to effectively inhibit tumor proliferation remains to be further explored through comprehensive functional studies.

### 3.3 Correlation Analysis Between Serine Metabolism Inhibition and Tumor Growth

Given that PSPH inhibitors Z1444603284, Z997780042, and Z1444669980, as well as PHGDH inhibitors NCT-503 and Compound 18, effectively reduced intracellular L-serine levels in tumor cell lines HCC-70 and BT-20, we further determine whether those inhibitors could inhibit tumor cells proliferation.

We first cultured HCC-70 and BT-20 cells in serine- and glycine-free DMEM medium and treated them with three PSPH inhibitors (Z1444603284, Z997780042, Z1444669980; concentration range: 0.156–40 μM) and two PHGDH inhibitors (NCT-503, Compound 18; concentration range: 0.078–20 μM) for 4 days. Cell proliferation was then assessed using the CCK-8 assay. The results showed that all three PSPH inhibitors-Z1444603284 (Fig. 3a–b), Z997780042 (Fig. 3c–d), and Z1444669980 (Fig. 3e–f) failed to significantly inhibit cell proliferation at any tested concentration, suggesting that although they reduced intracellular serine levels, this was insufficient to impair the proliferative capacity of tumor cells. In contrast, PHGDH inhibitors exhibited robust anti-proliferative effects. NCT-503 showed IC_50_ values of 18.2 μM and 10.4 μM for HCC-70 cells and BT-20 cells, respectively (Fig. 3g–h).Compound 18 exhibited a stronger potency with IC_50_ values of 6.0 μM and 5.9 μM (Fig. 3i–j), when compared to NCT-503. To determine whether the antiproliferative effects of PHGDH inhibitors were exclusively due to serine depletion, we repeated the above treatments in complete medium containing exogenous serine and glycine. We found that serine supplementation modestly increased the IC_50_ values of NCT-503 on HCC-70 cells from 18.2 μM to 28.2 μM and on BT-20 from 10.4 μM to 18.4 μM (Fig. 3g–h), and increased the IC_50_ values of Compound 18 on HCC-70 cells from 6.0 μM to 7.6 μM and on BT-20 from 5.9 μM to 10.6μM (Fig. 3i–j). The findings indicate that PHGDH inhibitors continue to demonstrate antiproliferative effects, even in the presence of exogenous supplementation with serine and glycine. These findings indicate that the anticancer effects of PHGDH inhibitors may not be solely dependent on serine deprivation but could involve additional metabolic pathways.

**Figure 3.**
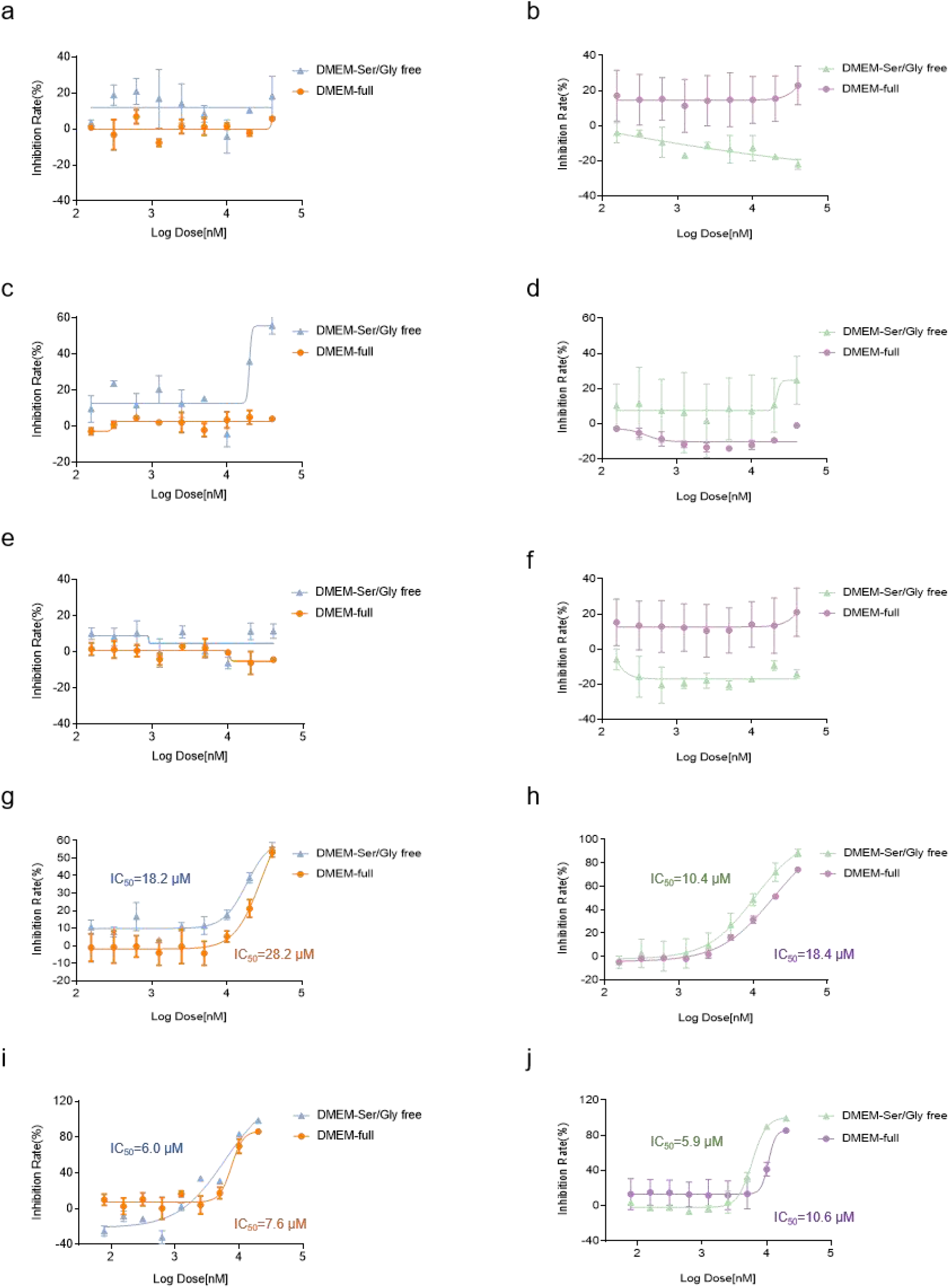
Correlation Analysis Between Serine Metabolism Inhibition and Tumor Growth. HCC-70 (left) and BT-20 (right) tumor cells were cultured in DMEM medium lacking serine and glycine, and treated with three PSPH inhibitors (Z1444603284, Z997780042, Z1444669980; concentration range: 0.156–40 μM) and two PHGDH inhibitors (NCT-503, Compound 18; concentration range: 0.078–20 μM) for 4 days. Cell viability was assessed using the CCK-8 assay. To further determine the role of serine in tumor cell proliferation, the same drug treatments were repeated in complete medium supplemented with exogenous serine and glycine for comparison. The resulting inhibition curves under both conditions are shown for Z1444603284 (a–b), Z997780042 (c–d), Z1444669980 (e–f), NCT-503 (g–h), and Compound 18 (i–j).The inhibition rate (Inh%) was calculated using the following formula: Inh% = 100 - [(RLU compound - RLU blank)/(RLU control - RLU blank)] × 100%.All data are presented as mean ± standard deviation (SD) (n = 6), and IC_50_ values were determined by nonlinear four-parameter logistic regression fitting.

In summary, this study compared the impact of PSPH and PHGDH inhibitors on serine metabolism and tumor cell proliferation. We demonstrate that inhibiting PHGDH exhibits greater potential as an anticancer strategy, primarily due to its broad impact on multiple metabolic pathways, rather than merely suppressing serine metabolism.

### 3.4 α-KG partially reversed the antitumor effect of PHGDH inhibition

As the upstream rate-limiting enzyme in the serine biosynthesis pathway, PHGDH not only catalyzes the conversion of 3-phosphoglycerate (3PG) to 3-phosphohydroxypyruvate (3PHP), but also produces two critical metabolic byproducts NADH and α-KG (Fig.4a). NADH acts as a key redox cofactor involved in the electron transport chain and oxidative phosphorylation. α-KG, an important intermediate of the tricarboxylic acid (TCA) cycle, affects cell proliferation by regulating energy metabolism, amino acid synthesis, one-carbon metabolism and lipid biosynthesis. Previous studies have shown that PHGDH knockdown reduces intracellular α-ketoglutarate (α-KG) levels by approximately 20%, suggesting that PHGDH inhibition may suppress tumor growth by impairing α-KG-mediated metabolic pathways[17].

To further investigate whether the anti-proliferative effects of PHGDH inhibition depend on α-KG, we treated HCC-70 and BT-20 tumor cells with PHGDH inhibitors NCT-503 and Compound 18 in serine/glycine-free DMEM medium, while supplementing with exogenous α-KG? After 4 days of treatment, cell proliferation was assessed using the CCK-8 assay. Results showed that α-KG supplementation partially rescued cell proliferation under PHGDH inhibition. In HCC-70 cells, the IC_50_ of NCT-503 increased from 18.2 μM to 24.5 μM (Fig. 4b), and the IC_50_ of Compound 18 increased slightly from 6.0 μM to 7.7 μM (Fig. 4d). In BT-20 cells, this effect was more pronounced, with the IC_50_ of NCT-503 increasing from 10.4 μM to 15.8 μM (Fig. 4c), and that of Compound 18 from 5.9 μM to 11.9 μM (Fig. 4e).

**Figure 4.**
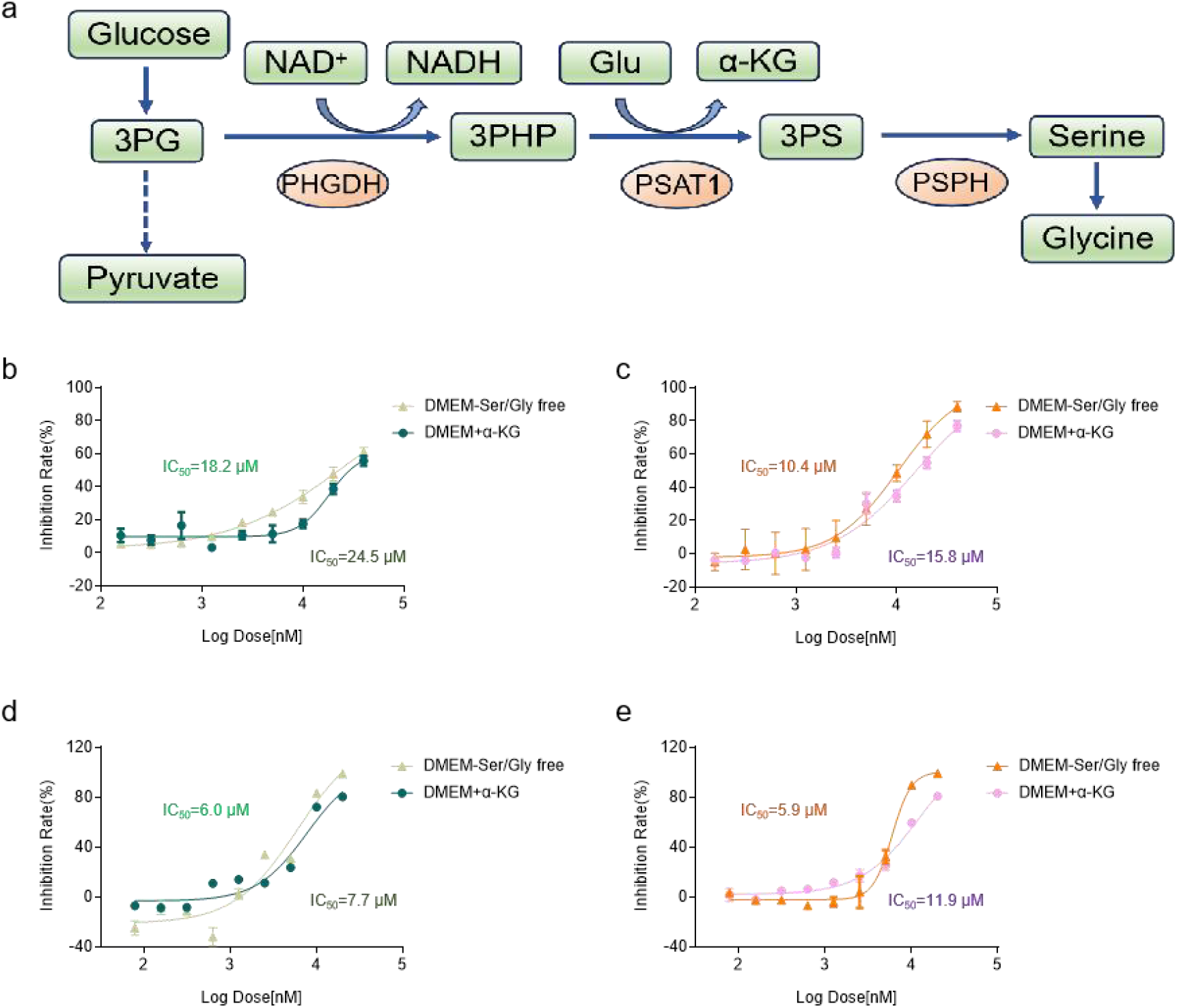
α-KG Rescue Reveals Non-Serine Roles of PHGDH in Tumor Metabolism. (a) Schematic diagram of the major pathway of serine metabolism. HCC-70 (left) and BT-20 (right) tumor cells were cultured in DMEM medium lacking serine and glycine, and treated with two PHGDH inhibitors (NCT-503 and Compound 18; concentration range: 0.078–20 μM) for 4 days. To further evaluate the role of α-ketoglutarate in tumor cell proliferation, an additional group was supplemented with α-ketoglutarate (final concentration: 1 mM) under identical treatment conditions. Dose–response curves of NCT-503 (b–c) and Compound 18 (d–e) in the absence or presence of α-ketoglutarate are shown. Inhibition rate (Inh%) was calculated as: Inh% = 100 − [(RLU compound − RLU blank) / (RLU control − RLU blank)] × 100%. Data are presented as mean ± standard deviation (SD) (n = 6). IC_50_ values were determined by nonlinear four-parameter logistic regression.

Collectively, these results indicate that the antitumor effects of PHGDH inhibition are not solely attributable to suppression of serine biosynthesis, but instead result from coordinated disruption of multiple metabolic pathways-including the TCA cycle, one-carbon metabolism, and redox balance-that together destabilize cellular metabolic homeostasis and suppress tumor cell proliferation. This finding challenges the oversimplified view of “serine as the sole target” and highlights the broader regulatory role of PHGDH in metabolic networks.

## 4 Discussion

Metabolic reprogramming in tumor cells has emerged as a central theme in recent cancer research. Among various metabolic pathways, serine metabolism is recognized as a critical anabolic route that supports rapid cell proliferation and division in cancer cells[18].Targeting the serine biosynthesis pathway has thus become a promising anticancer strategy; however, the exact mechanisms by which inhibition of serine synthesis limits tumor growth remain incompletely understood.

PHGDH, the upstream rate-limiting enzyme of this pathway, is frequently overexpressed in multiple cancer types, making it a key target for metabolic intervention[19]. In this study, we systematically compared the effects of targeting PHGDH and PSPH-two distinct enzymatic nodes in the serine pathway-on serine levels and proliferative capacity in tumor cells. Through high-throughput screening and molecular docking, we developed and validated three novel PSPH inhibitors that effectively reduced intracellular L-serine levels across various cancer cell lines. However, unlike PHGDH inhibitors, these PSPH inhibitors failed to significantly suppress tumor cell proliferation, suggesting that depletion of serine alone is insufficient to block tumor growth. Furthermore, even in the presence of exogenous serine supplementation, PHGDH inhibitors continued to exhibit robust antiproliferative effects, implying that their anticancer activity is not entirely dependent on serine deprivation.Notably, previous studies have shown that under serine-limiting conditions, the normal sphingolipid synthesis pathway can shift toward the production of toxic deoxysphingolipids, which exert cytotoxic effects on tumor cells. In tumor-bearing mice, treatment with the PHGDH inhibitor PH-755 for one week resulted in significant tumor shrinkage, along with disruptions in sphingolipid and deoxysphingolipid balance, thereby impeding tumor progression[20].

From a metabolic perspective, PHGDH catalyzes the conversion of 3-phosphoglycerate (3PG) to 3-phosphohydroxypyruvate (3PHP), during which NAD^+^ is reduced to NADH and glutamate is deaminated to form α-ketoglutarate (α-KG). These byproducts play central roles in multiple critical metabolic networks. We experimentally confirmed one of these mechanisms by supplementing exogenous α-KG, which partially reversed the antiproliferative effects of PHGDH inhibition. Beyond its role as a core intermediate in the tricarboxylic acid (TCA) cycle[21], α-KG is involved in glutamine metabolism, lipid biosynthesis, and epigenetic regulation. Studies have indicated that dysregulated glutamine metabolism is essential for tumorigenesis, microenvironment remodeling, and therapeutic resistance[22]. As a key glutamine-derived metabolite, α-KG accumulation exerts tumor-suppressive effects by redirecting glucose metabolism[23], promoting cancer cell differentiation[24], and triggering oxidative stress-induced ferroptosis[25]. Therefore, the depletion of α-KG caused by PHGDH inhibition may disrupt carbon metabolism and redox balance, thereby impairing tumor adaptive growth.

In addition, PHGDH inhibition reduces the production of NADH, a key electron donor in redox reactions. NADH primarily fuels oxidative phosphorylation in the mitochondrial electron transport chain, generating ATP to support cancer cell growth[26]. A decrease in NADH levels may reduce energy production and lead to accumulation of reactive oxygen species (ROS), initiating oxidative stress, mitochondrial dysfunction, and subsequent apoptosis or senescence. Changes in the NADH/NAD^+^ ratio can also influence the activity of NAD^+^-dependent enzymes such as Sirtuins and PARPs, thereby affecting gene expression and DNA repair processes[27].These observations warrant further investigation.

We speculate that PHGDH inhibitors exert anticancer effects through multi-target, multi-pathway metabolic disruption, including inhibition of serine synthesis, suppression of α-KG-related pathways, and disturbance of NADH-mediated redox balance. These combined metabolic pressures effectively compromise tumor cell survival and proliferation. However, the systemic nature of these metabolic disruptions also raises concerns regarding safety. For instance, PHGDH knockout mice exhibit severe central nervous system malformations and embryonic lethality[28], while treatment with NCT-503 has been reported to halt embryonic development, likely due to its ability to cross the blood-brain barrier[29].

In conclusion, the antitumor mechanisms of PHGDH inhibitors extend far beyond serine suppression alone. Their broad, multi-dimensional metabolic interventions offer potent and systematic therapeutic potential. Nonetheless, achieving a balance between efficacy and safety remains a critical challenge, underscoring the need for further research to guide the development of more precise and safer metabolic-targeted cancer therapies.

## Supporting information

Analysis of Compound Purity

## Abbreviations

3PG: 3-phosphoglycerate
3PHP: 3-phosphohydroxypyruvate
PHGDH: phosphoglycerate dehydrogenase
PSAT1: phosphoserine aminotransferase 1
p-Ser: phosphoserine
α-KG: α-ketoglutarate
PSPH: phosphoserine phosphatase
NADH: Nicotinamide adenine dinucleotide
SPR: Surface plasmon resonance
HTVS: high-throughput virtual screening
Kd: dissociation constants
BBB: blood-brain barrier
CNS: central nervous system
TCA: tricarboxylic acid
ROS: reactive oxygen species
ATP: Adenosine 5’-triphosphate
PTM: spost-translational modifications

## CRediT authorship contribution statement

Wang Yanbing: Writing-review & editing, Writing-original draft, Formal analysis, Data curation.Sha Longze:Writing -review & editing, Project administration, Methodology, Funding acquisition, Conceptualization.

## Data availability statement

Source data are provided with this paper. All other data are available from the corresponding authors upon reasonable request.

## Acknowledgement

None.

---

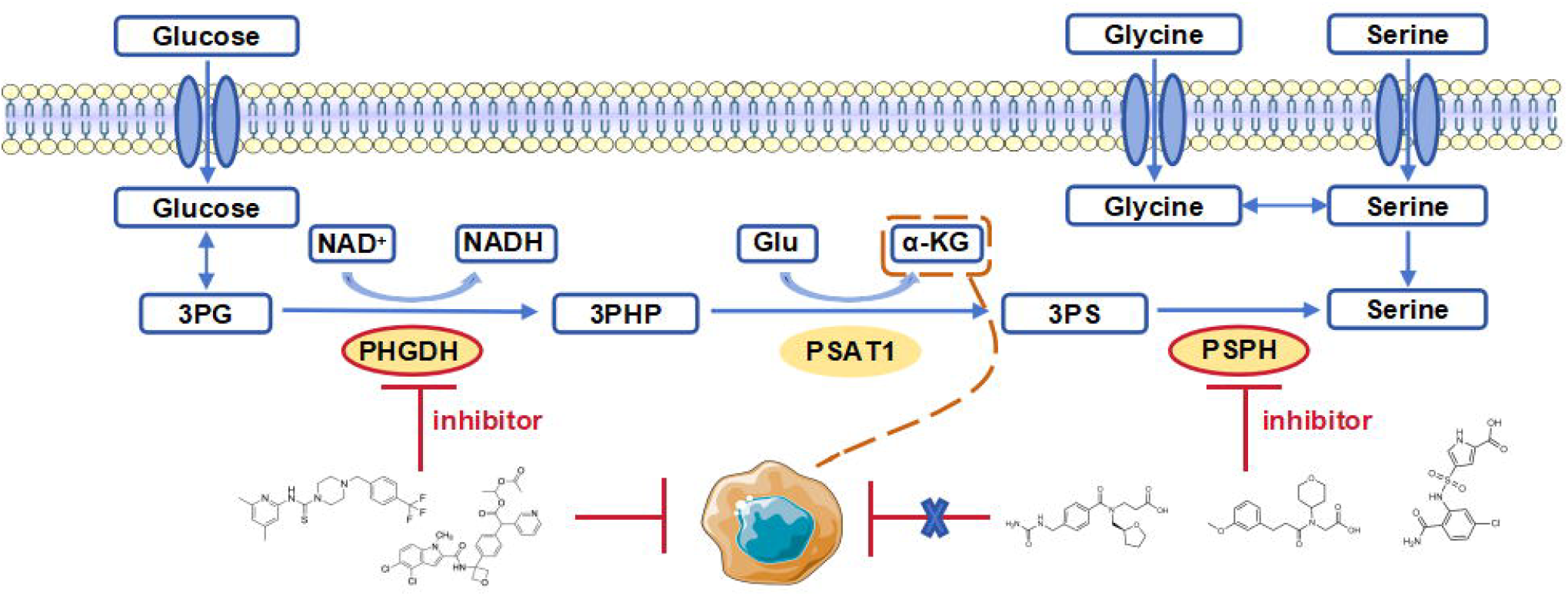

## References

1. Organization, W.H., Global cancer burden growing, amidst mounting need for services. World Health Organization: Geneva, Switzerland, 2024.

2. McNamee, M.J., D. Michod, and M.V. Niklison-Chirou, Can small molecular inhibitors that stop de novo serine synthesis be used in cancer treatment? Cell Death Discovery, 2021. 7(1): p. 87.

3. Geeraerts, S.L., et al., The ins and outs of serine and glycine metabolism in cancer. Nature Metabolism, 2021. 3(2): p. 131–141.

4. Pavlova, N.N. and C.B. Thompson, The Emerging Hallmarks of Cancer Metabolism. Cell Metab, 2016. 23(1): p. 27–47.

5. Yang, M. and K.H. Vousden, Serine and one-carbon metabolism in cancer. Nature Reviews Cancer, 2016. 16(10): p. 650–662.

6. Jain, M., et al., Metabolite profiling identifies a key role for glycine in rapid cancer cell proliferation. Science, 2012. 336(6084): p. 1040–4.

7. Shunxi, W., et al., Serine Metabolic Reprogramming in Tumorigenesis, Tumor Immunity, and Clinical Treatment. Advances in Nutrition, 2023. 14(5): p. 1050–1066.

8. Possemato, R., et al., Functional genomics reveal that the serine synthesis pathway is essential in breast cancer. Nature, 2011. 476(7360): p. 346–50.

9. Zhang, B., et al., PHGDH Defines a Metabolic Subtype in Lung Adenocarcinomas with Poor Prognosis. Cell Rep, 2017. 19(11): p. 2289–2303.

10. Zhu, J., et al., High Expression of PHGDH Predicts Poor Prognosis in Non-Small Cell Lung Cancer. Transl Oncol, 2016. 9(6): p. 592–599.

11. Li, A.M. and J. Ye, The PHGDH enigma: Do cancer cells only need serine or also a redox modulator? Cancer Lett, 2020. 476: p. 97–105.

12. Lee, C.M., et al., PHGDH: a novel therapeutic target in cancer. Exp Mol Med, 2024. 56(7): p. 1513–1522.

13. Chen, J., et al., Phosphoglycerate dehydrogenase is dispensable for breast tumor maintenance and growth. Oncotarget, 2013. 4(12): p. 2502–11.

14. Mullarky, E., et al., Inhibition of 3-phosphoglycerate dehydrogenase (PHGDH) by indole amides abrogates de novo serine synthesis in cancer cells. Bioorg Med Chem Lett, 2019. 29(17): p. 2503–2510.

15. Pacold, M.E., et al., A PHGDH inhibitor reveals coordination of serine synthesis and one-carbon unit fate. Nature Chemical Biology, 2016. 12(6): p. 452–458.

16. Chen, L., S. Liu, and Y. Tao, Regulating tumor suppressor genes: post-translational modifications. Signal Transduction and Targeted Therapy, 2020. 5(1): p. 90.

17. Wei, L., et al., Genome-wide CRISPR/Cas9 library screening identified PHGDH as a critical driver for Sorafenib resistance in HCC. Nature Communications, 2019. 10(1): p. 4681.

18. Maddocks, O.D., et al., Serine Metabolism Supports the Methionine Cycle and DNA/RNA Methylation through De Novo ATP Synthesis in Cancer Cells. Mol Cell, 2016. 61(2): p. 210–21.

19. Birsoy, K., L.A. Garraway, and M. Mino-Kenudson, Functional genomics reveals serine synthesis is essential in PHGDH-amplified breast cancer. Nature, 2012. 476(7360): p. 346–350.

20. Muthusamy, T., et al., Serine restriction alters sphingolipid diversity to constrain tumour growth. Nature, 2020. 586(7831): p. 790–795.

21. Cai, Y., et al., α-KG inhibits tumor growth of diffuse large B-cell lymphoma by inducing ROS and TP53-mediated ferroptosis. Cell Death Discov, 2023. 9(1): p. 182.

22. Yang, L., S. Venneti, and D. Nagrath, Glutaminolysis: A Hallmark of Cancer Metabolism. Annu Rev Biomed Eng, 2017. 19: p. 163–194.

23. Zhang, J.Y., et al., The metabolite α-KG induces GSDMC-dependent pyroptosis through death receptor 6-activated caspase-8. Cell Res, 2021. 31(9): p. 980–997.

24. Morris, J.P.t., et al., α-Ketoglutarate links p53 to cell fate during tumour suppression. Nature, 2019. 573(7775): p. 595–599.

25. Shin, D., et al., Dihydrolipoamide dehydrogenase regulates cystine deprivation-induced ferroptosis in head and neck cancer. Redox Biol, 2020. 30: p. 101418.

26. Cairns, R.A., I.S. Harris, and T.W. Mak, Regulation of cancer cell metabolism. Nat Rev Cancer, 2011. 11(2): p. 85–95.

27. Imai, S. and L. Guarente, NAD+ and sirtuins in aging and disease. Trends Cell Biol, 2014. 24(8): p. 464–71.

28. Furuya, S., An essential role for de novo biosynthesis of L-serine in CNS development. Asia Pac J Clin Nutr, 2008. 17 Suppl 1: p. 312–5.

29. Yoshida, K., et al., Targeted disruption of the mouse 3-phosphoglycerate dehydrogenase gene causes severe neurodevelopmental defects and results in embryonic lethality. J Biol Chem, 2004. 279(5): p. 3573–7.

