## Supplementary material for "Comparative Analysis of the Effects of PSPH and PHGDH Inhibitors on Tumor Cell Proliferation": Analysis of Compound Purity

**Supplementary Note**

The purity of the four PSPH small-molecule inhibitors was evaluated by mass spectrometry, and the representative spectra are presented below.

**1.** Mass spectrometric analysis of Z218484536

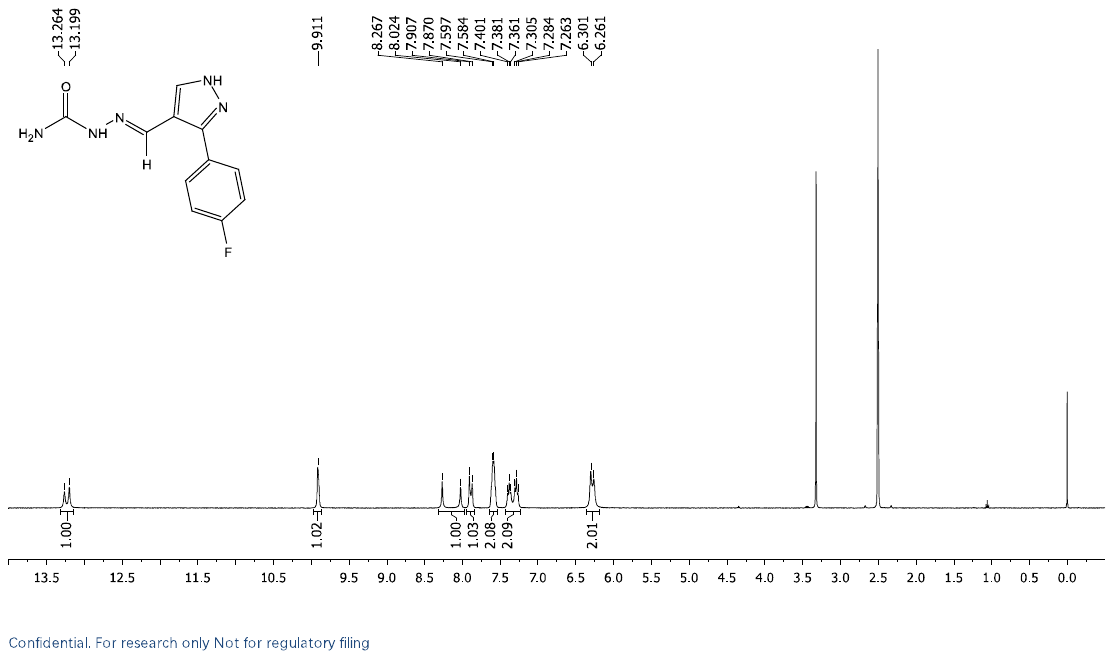

Figure 1. Mass spectrometric analysis of Z218484536.

**2.** Mass spectrometric analysis of Z1444669980

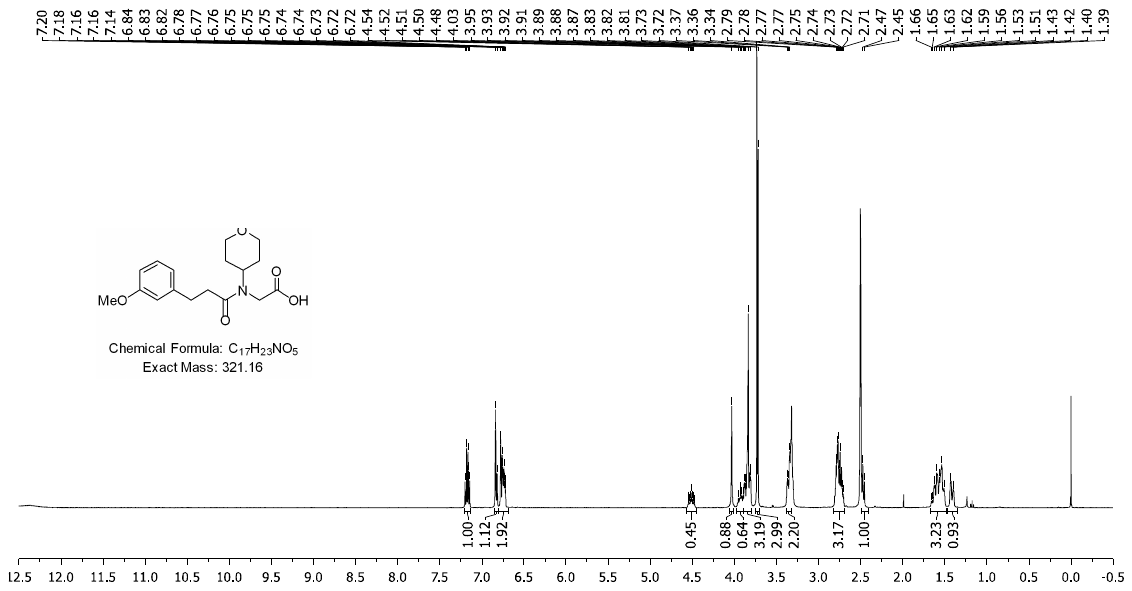

Figure 2. Mass spectrometric analysis of Z1444669980.

**3.** Mass spectrometric analysis of Z1444603284

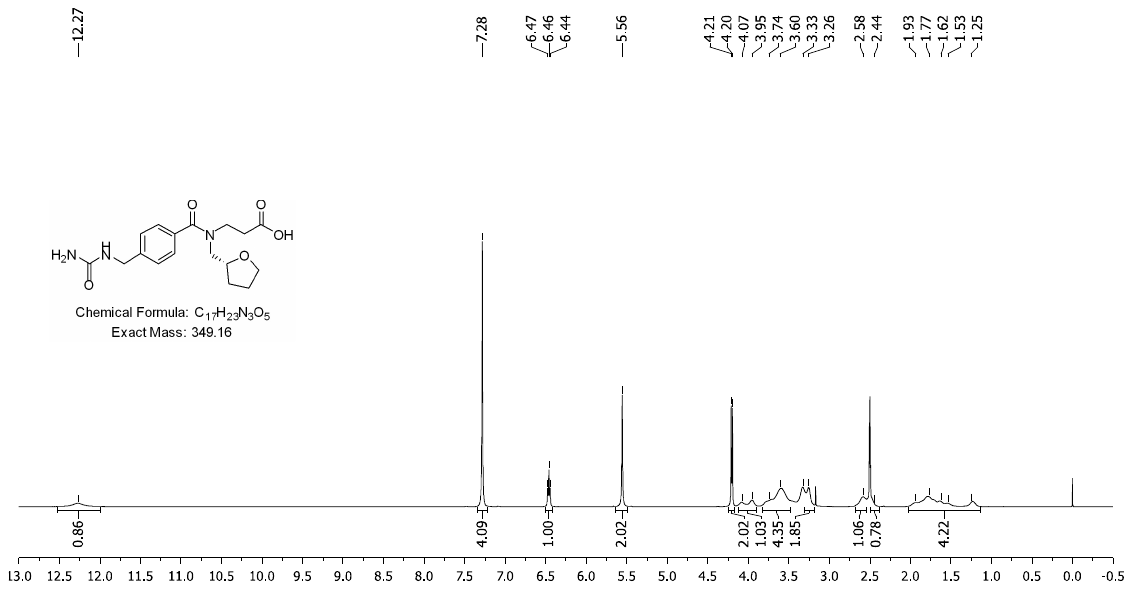

Figure 3. Mass spectrometric analysis of Z1444603284.

**4.** Mass spectrometric analysis of Z997780042

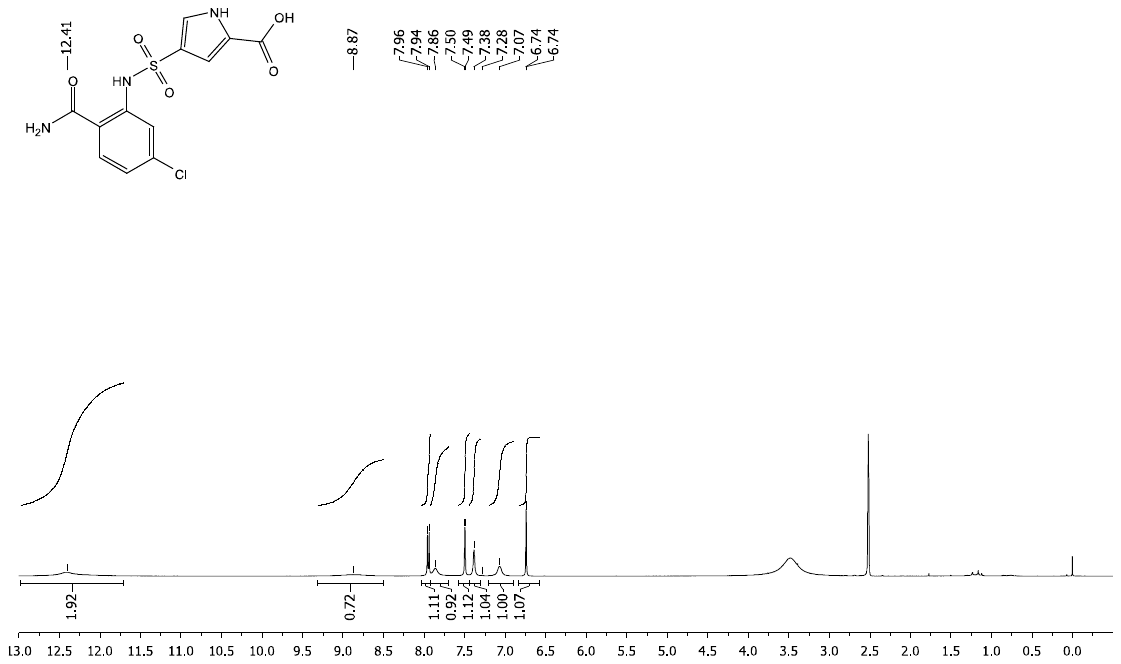

Figure 4. Mass spectrometric analysis of Z997780042.
